# HoCoRT: Host contamination removal tool

**DOI:** 10.1101/2022.11.18.517030

**Authors:** Ignas Rumbavicius, Trine B. Rounge, Torbjørn Rognes

## Abstract

**Background:** Shotgun metagenome sequencing data obtained from a host environment will usually be contaminated with sequences from the host organism. Host sequences should be removed before further analysis to avoid biases, to reduce downstream computational load, or for privacy reasons in case of a human host. The tools that we identified as designed specifically to perform host contamination sequence removal were either outdated, not maintained or complicated to use. We have therefore created HoCoRT, a fast and user friendly tool that implements several methods for optimised host sequence removal, and evaluated the speed and accuracy of these methods.

**Results:** HoCoRT is an open-source command-line tool for host contamination removal. It is designed to be simple to install and use, including a one-step option for genome indexing. HoCoRT uses a range of well-known mapping, classification and alignment methods for the classification of reads. The underlying classification method to be used and its parameters are selectable by the user, enabling adaptation to different circumstances. Based on our investigation of the performance of a range of methods and parameters on artificial human gut and oral microbiomes, recommendations are provided for typical data sets with short and long reads.

**Conclusions:** To decontaminate a human microbiome with HoCoRT, the best combination of speed and accuracy was found with BioBloom, Bowtie2 in end-to-end mode, or HISAT2 for short reads, while BioBloom performed best for long reads. For the oral microbiome, Bowtie2 was slightly more accurate than the other two, at the expense of speed. HoCoRT using Bowtie2 in end-to-end mode was found to be much faster and slightly more accurate than the dedicated DeconSeq tool. HoCoRT is distributed as a Bioconda package. Source code and documentation are available at https://github.com/ignasrum/hocort for Linux and macOS under the MIT licence.

## Background

Sequencing the genomes of microbial communities in an environment of a host organism have opened new avenues for research on host - microbe interactions. A number of analysis steps must be carried out after metagenomic sequencing to obtain a good picture of the composition of the microbiome. The huge amounts of data often involved require efficient processing, and it is therefore important to avoid unnecessary computations. Some sequences of host origin are inevitably included in the obtained data, and when the host is human, privacy becomes an issue. Sequences of non-microbial origin could also bias the results of downstream analyses. These host sequences should hence be removed at an early stage [1].

Decontamination is often handled in an ad-hoc manner by running an alignment tool to search reads against a host genome database. This makes this essential analysis step more complicated than necessary and could result in inferior performance. There is a lack of best practices for this procedure, and comparison across studies is more difficult without standardisation.

We could identify only two dedicated tools specifically designed for removal of contaminating sequences, namely DeconSeq and GenCoF. DeconSeq [2] is a command-line tool for identification and removal of sequence contamination from genomic and metagenomic datasets. DeconSeq integrates its underlying classifier, a modified version of BWA-SW [3], directly into its source code, making modifications difficult. DeconSeq only supports single-end Illumina reads, and the code has not been updated since 2013. GenCoF [4] is a graphical user interface to rapidly remove human genome contaminants from metagenomic datasets. It is limited to short reads. The GUI makes it unsuited for scripting. Extending GenCoF is also difficult as it also has its underlying classifier Bowtie2 [5] directly integrated into its source code. Both tools clearly have severe limitations making them less suitable for most large-scale modern datasets. We therefore decided to develop the new tool HoCoRT to fill this need. We also examined the performance of a range of underlying classification methods in order to be able to make recommendations about which method to use under different circumstances and to provide default settings.

## Implementation

HoCoRT is an open-source command-line based tool written in Python 3. It is designed to be easy to use and can be easily installed as a package with Bioconda [6], or using a Docker container. HoCoRT has a modular pipeline design and uses well established classification, mapping and alignment tools to perform the classification of the sequences into host and non-host (microbial) sequences. Currently available pipeline modules include the BBMap tool in the BBTools suite [7, 8], as well as BioBloom [9], Bowtie2 [5], BWA-MEM2 [10], HISAT2 [11], Kraken2 [12], and Minimap2 [13]. Furthermore, modules may pipe data through different tools after each other. The configuration of the options to the pipelines can be specified by the user, but recommended defaults are provided. New modules can be written to extend HoCoRT to use other tools. HoCoRT also provides an extensive Python library with an API that could be used as a backend by other tools. HoCoRT reads and writes optionally compressed FASTQ files. The building of database index files for the underlying classification tools is also handled by HoCoRT. The tool is documented with built-in help functions and error messages. HoCoRT depends on Samtools [14]. Please see table 1 for a list of all software mentioned in this work, including version numbers and references.

**Table 1.** Software. The software packages used or mentioned are listed with version numbers and references.

| Software | Version | References |
| --- | --- | --- |
| BBDuk | 39.01 | [7, 8] |
| BBMap | 39.01 | [7, 8] |
| BBSplit | 39.01 | [7, 8] |
| BioBloom Tools | 2.3.5 | [9] |
| BioConda | 23.1.0 | [6] |
| Bowtie2 | 2.4.5 | [5] |
| BWA-SW |  | [3] |
| BWA-MEM2 | 2.2.1 | [10] |
| CONSULT |  | [18] |
| DeconSeq | 0.4.3 | [2] |
| GenCoF |  | [4] |
| HISAT2 | 2.2.1 | [11] |
| HoCoRT | 1.2.2 | [20] |
| InSiliconSeq | 1.5.4 | [16] |
| Kraken2 | 2.1.2 | [12] |
| MiniMap2 | 2.24 | [13] |
| NanoSim | 3.0.0 | [17] |
| Samtools | 1.16.1 | [14] |
| Seal | 39.01 | [7, 8] |
| SnakeMake | 7.19.1 | [19] |

## Results and discussion

The classification speed and accuracy of HoCoRT with several different underlying methods and settings was investigated with synthetic datasets. The GitHub repository at https://github.com/ignasrum/hocort-eval provides the scripts used for generating the datasets and to perform the performance evaluation. The datasets were created with HiSeq, MiSeq and Nanopore reads containing either a mix of 1% human host sequences and 99% microbial sequences, similar to a human gut microbiome, or a mix of 50% human host and 50% microbial sequences, similar to a human oral microbiome.

The human reads were generated from the Genome Reference Consortium Human Build 38 patch release 13 (GRCh38.p13), whereas the microbial reads are generated from a mix of bacterial, fungal and viral sequences pseudo-randomly extracted from NCBI GenBank [15]. HoCoRT was run using several different pipelines (classification modules) and parameters on HiSeq, MiSeq and Nanopore data. For each of these combinations, 7 different datasets were analysed, each consisting of 5 million reads randomly generated using InSilicoSeq [16] (HiSeq 125bp and MiSeq 300bp, paired-end) and NanoSim [17] (Nanopore, average 2159bp, range 54-98320bp, single-end).

For Illumina data the following 17 pipelines were examined: Seal, BBduk, BBsplit, BioBloom, Bowtie2 in end-to-end and local mode, both with and without the “un_conc” option, HISAT2, Kraken2, BBMap in default and fast mode, BWA-MEM2, Kraken2 followed by Bowtie2 in end-to-end mode, Kraken2 followed by HISAT2, Minimap2, and finally Kraken2 followed by Minimap2. For Nanopore data the following 4 pipelines were examined: BioBloom, Minimap2, Kraken2 followed by Minimap2, and Kraken2. When the “un_conc” option is given, Bowtie2 requires both reads in a pair to map concordantly to the genome. We also considered including CONSULT [18] in the comparison, but there were no pre-compiled binaries or packages available, and it requires 128GB of RAM, making it impractical for many users.

The ability to detect the human host sequences was tested and the sensitivity, precision, and accuracy was calculated, considering the correctly identified human host sequences as true positives. We considered accuracy as most important, followed by speed. Performance analysis was carried out using a Snakemake pipeline [19] on a desktop PC with an AMD Ryzen 7 1700X 8 core/16 thread 3.4 GHz CPU, 64GB RAM and 4TB HDD running Linux. No quality filtering or other pre-processing was performed.

The results for the gut microbiome are shown in figure 1, while additional results are included in Supplementary figure S1 and table S1. Results for the oral microbiome are shown in Supplementary figure S2 and table S2. Overall, BioBloom, Bowtie2 in end-to-end mode, and HISAT2 performed best for Illumina reads and are recommended as they are highly accurate in all cases and also generally very fast. Bowtie2 had the highest accuracy for the oral microbiome, at the expense of speed. For Nanopore reads, BioBloom performed best and is recommended due to its combination of high accuracy and speed.

**Fig. 1.**
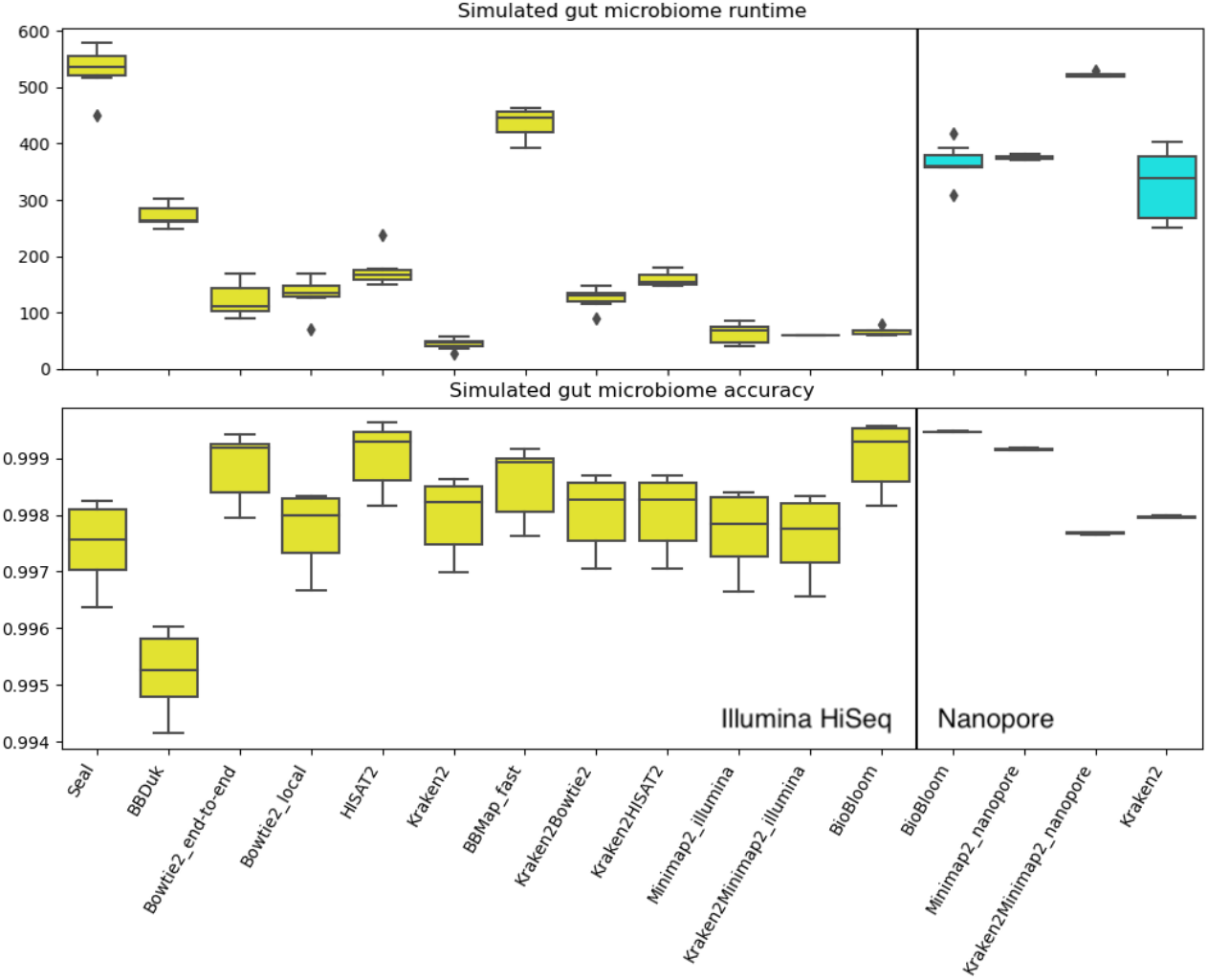
HoCoRT performance on simulated gut microbiome datasets. Boxplots of HoCoRT runtime in seconds (top) and classification accuracy (bottom) using several different classification modules and parameters on Illumina (HiSeq) (yellow, left), and Nanopore data (cyan, right). Supplementary figure S1 is similar, but also includes results for MiSeq data. Supplementary table S1 contains additional results, including those for BBMap in default mode, BBsplit, Bowtie2 with the “un_conc” option, and BWA-MEM2, which were excluded from the figures due to outliers.

The performance of HoCoRT was also compared to DeconSeq using a human gut dataset, but with single-ended Illumina reads. The HoCoRT Bowtie2 (end-to-end) pipeline was much faster than DeconSeq at aligning both HiSeq (34X) and MiSeq (49X) reads, and also slightly more accurate, as shown in Supplementary table S3.

Additional results and a more detailed description of HoCoRT can be found in the first author’s master thesis [20].

## Conclusions

A dedicated, flexible, extendable, and modular tool for host sequence contamination removal is now freely available for all users. A comparison of the available classification methods has been performed and recommendations are provided. The HoCoRT tool should simplify the decontamination step in microbiome data analysis and provide solid performance.

## Availability and requirements

**Project name:** HoCoRT

**Project home page:** https://github.com/ignasrum/hocort

**Operating system(s):** Linux and macOS

**Programming language:** Python

**Other requirements:** Samtools [14] and other packages. Please see GitHub repository for details.

**License:** MIT license

**Any restrictions to use by non-academics:** None

## Acknowledgements

Thanks to the anonymous reviewers of a previous version of the manuscript for suggesting additional tools to consider.

## Authors’ contributions

IR developed and evaluated the tool, performed experiments, made tables and figures and wrote the initial description of the tool. TR initiated the project and drafted the manuscript. TBR provided datasets. TBR and TR provided advice in all stages of the project. All authors analysed and interpreted the results. All authors edited, read and approved the final manuscript.

## Ethics declarations

### Ethics approval and consent to participate

Not applicable.

### Consent for publication

Not applicable.

