## Supplementary material for "HoCoRT: Host contamination removal tool"

#### Supplementary figures

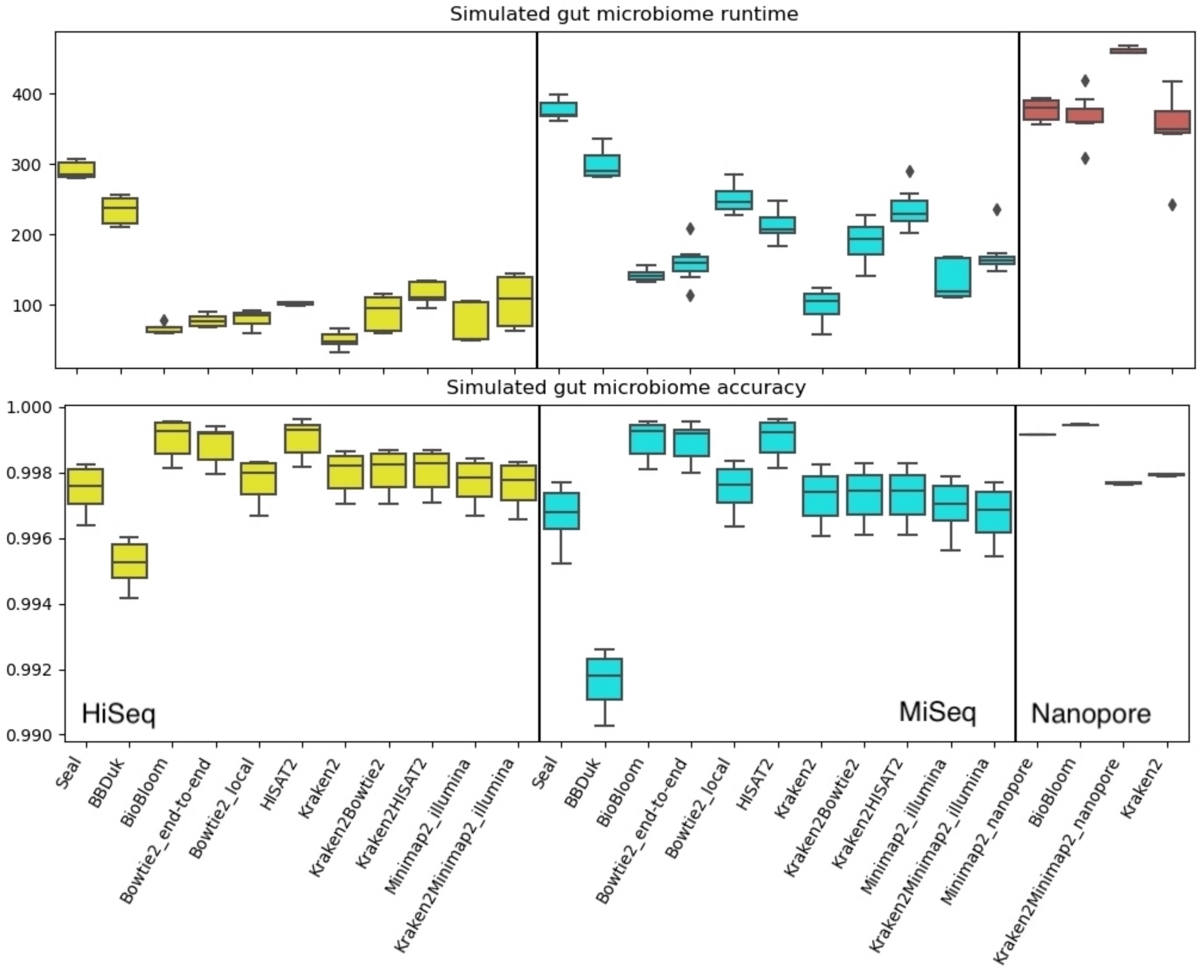


**Fig. S1. HoCoRT performance on simulated gut microbiome datasets**. Boxplots of HoCoRT runtime in seconds (top) and classification accuracy (bottom) using several different classification modules and parameters on HiSeq (yellow, left), MiSeq (cyan, middle) and Nanopore data (red, right). BBMap, BBSplit, Bowtie2 with the "un_conc" option, and BWA-MEM2 were excluded due to outliers.

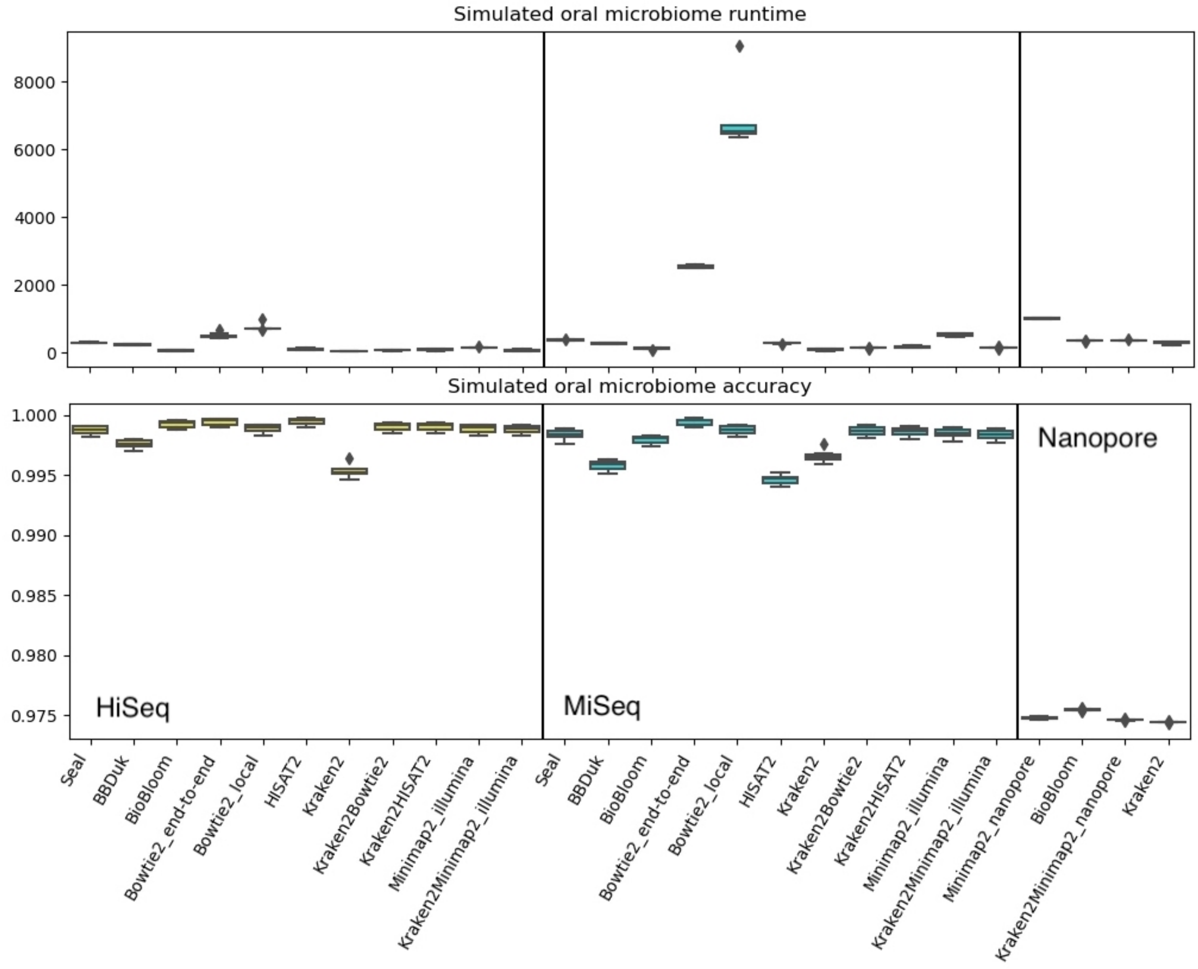


**Fig. S2. HoCoRT performance on simulated oral microbiome datasets.** Boxplots of HoCoRT runtime in seconds (top) and classification accuracy (bottom) using several different classification modules and parameters on HiSeq (yellow, left), MiSeq (cyan, middle) and Nanopore data (red, right). BBMap, BBSplit, Bowtie2 with the “un_conc” option, and BWA-MEM2 were excluded due to outliers.

### Supplementary tables

**Table S1. Detailed HoCORT performance on simulated gut microbiome datasets.** The average runtime (in seconds), accuracy, precision, and sensitivity is shown for each pipeline and for each data type. The best (blue) and worst (red) performing pipelines are indicated for each performance metric and data type.

| Pipeline | Runtime | Accuracy | Precision | Sensitivity |
|---|---|---|---|---|
| **Paired-end Hiseq** | | | | |
| Seal | 291.3 | 0.9975 | 0.8027 | 1.0000 |
| BBDuk | 233.7 | 0.9952 | 0.6786 | 1.0000 |
| BBSplit | 509.0 | 0.9982 | 0.8523 | 1.0000 |
| BioBloom | 66.6 | 0.9990 | 0.9143 | 0.9995 |
| Bowtie2_end-to-end | 77.4 | 0.9988 | 0.8978 | 1.0000 |
| Bowtie2_local | 80.4 | 0.9978 | 0.8187 | 1.0000 |
| Bowtie2_end-to-end_un_conc | 277.2 | 0.9934 | 0.9351 | 0.3625 |
| Bowtie2_local_un_conc | 314.9 | 0.9941 | 0.8956 | 0.4614 |
| HISAT2 | 101.7 | 0.9990 | 0.9145 | 0.9998 |
| Kraken2 | 49.8 | 0.9980 | 0.8385 | 0.9928 |
| BBMap_default | 1053.2 | 0.9982 | 0.8520 | 1.0000 |
| BBMap_fast | 300.9 | 0.9986 | 0.8762 | 0.9999 |
| BWA_MEM2 | 381.3 | 0.9720 | 0.2635 | 1.0000 |
| Kraken2Bowtie2 | 87.7 | 0.9980 | 0.8385 | 1.0000 |
| Kraken2HISAT2 | 117.2 | 0.9980 | 0.8388 | 1.0000 |
| Minimap2_illumina | 73.3 | 0.9977 | 0.8170 | 1.0000 |
| Kraken2Minimap2_illumina | 105.2 | 0.9976 | 0.8107 | 1.0000 |
| **Paired-end MiSeq** | | | | |
| Seal | 376.7 | 0.9967 | 0.7559 | 1.0000 |
| BBDuk | 299.7 | 0.9916 | 0.5457 | 1.0000 |
| BBSplit | 791.9 | 0.9985 | 0.8726 | 1.0000 |
| BioBloom | 142.0 | 0.9990 | 0.9129 | 0.9969 |
| Bowtie2_end-to-end | 159.0 | 0.9989 | 0.9041 | 0.9999 |
| Bowtie2_local | 249.8 | 0.9975 | 0.8043 | 1.0000 |
| Bowtie2_end-to-end_un_conc | 747.3 | 0.9904 | 0.9721 | 0.0457 |
| Bowtie2_local_un_conc | 810.6 | 0.9919 | 0.8761 | 0.2243 |
| HISAT2 | 212.6 | 0.9990 | 0.9224 | 0.9901 |
| Kraken2 | 99.0 | 0.9973 | 0.7902 | 0.9960 |
| BBMap_default | 2338.7 | 0.9985 | 0.8730 | 0.9993 |
| BBMap_fast | 733.3 | 0.9989 | 0.9044 | 0.9956 |
| BWA_MEM2 | 2889.4 | 0.9128 | 0.1032 | 1.0000 |
| Kraken2Bowtie2 | 189.2 | 0.9973 | 0.7908 | 1.0000 |
| Kraken2HISAT2 | 236.2 | 0.9973 | 0.7908 | 1.0000 |
| Minimap2_illumina | 136.5 | 0.9970 | 0.7698 | 1.0000 |
| Kraken2Minimap2_illumina | 170.9 | 0.9967 | 0.7567 | 1.0000 |
| **Single-end Nanopore** | | | | |
| BioBloom | 366.0 | 0.9995 | 0.9957 | 0.9506 |
| Minimap2_nanopore | 376.3 | 0.9992 | 0.9660 | 0.9494 |
| KrakenMinimap2_nanopore | 460.6 | 0.9977 | 0.8386 | 0.9507 |
| Kraken2 | 349.3 | 0.9979 | 0.8591 | 0.9499 |

**Table S2. Detailed HoCoRT performance on simulated oral microbiome datasets.** The average runtime (in seconds), accuracy, precision, and sensitivity of the classification is shown for each pipeline and for each data type. The best (blue) and worst (red) performing pipelines are indicated for each performance metric and data type.

| Pipeline | Runtime | Accuracy | Precision | Sensitivity |
|---|---|---|---|---|
| **Paired-end Hiseq** | | | | |
| Seal | 289.5 | 0.9987 | 0.9975 | 1.0000 |
| BBDuk | 228.2 | 0.9976 | 0.9952 | 1.0000 |
| BBSplit | 1489.1 | 0.9991 | 0.9982 | 1.0000 |
| BioBloom | 60.0 | 0.9992 | 0.9990 | 0.9994 |
| Bowtie2_end-to-end | 495.4 | 0.9994 | 0.9988 | 1.0000 |
| Bowtie2_local | 739.4 | 0.9989 | 0.9977 | 1.0000 |
| Bowtie2_end-to-end_un_conc | 571.3 | 0.6809 | 0.9993 | 0.3621 |
| Bowtie2_local_un_conc | 814.7 | 0.7300 | 0.9988 | 0.4606 |
| HISAT2 | 104.4 | 0.9994 | 0.9990 | 0.9998 |
| Kraken2 | 47.5 | 0.9954 | 0.9980 | 0.9927 |
| BBMap_default | 9351.0 | 0.9991 | 0.9982 | 1.0000 |
| BBMap_fast | 1062.5 | 0.9992 | 0.9985 | 0.9999 |
| BWA_MEM2 | 461.9 | 0.9858 | 0.9725 | 1.0000 |
| Kraken2Bowtie2 | 68.1 | 0.9990 | 0.9980 | 1.0000 |
| Kraken2HISAT2 | 79.9 | 0.9990 | 0.9980 | 1.0000 |
| Minimap2_illumina | 149.1 | 0.9988 | 0.9977 | 1.0000 |
| Kraken2Minimap2_illumina | 70.9 | 0.9988 | 0.9976 | 1.0000 |
| **Paired-end MiSeq** | | | | |
| Seal | 374.8 | 0.9983 | 0.9967 | 1.0000 |
| BBDuk | 273.0 | 0.9958 | 0.9916 | 1.0000 |
| BBSplit | 4046.9 | 0.9993 | 0.9985 | 1.0000 |
| BioBloom | 116.9 | 0.9979 | 0.9990 | 0.9968 |
| Bowtie2_end-to-end | 2540.8 | 0.9994 | 0.9989 | 0.9999 |
| Bowtie2_local | 6887.7 | 0.9988 | 0.9975 | 1.0000 |
| Bowtie2_end-to-end_un_conc | 2534.2 | 0.5230 | 0.9997 | 0.0460 |
| Bowtie2_local_un_conc | 7015.4 | 0.6122 | 0.9986 | 0.2247 |
| HISAT2 | 267.9 | 0.9946 | 0.9991 | 0.9901 |
| Kraken2 | 89.4 | 0.9966 | 0.9973 | 0.9959 |
| BBMap_default | 84978.6 | 0.9989 | 0.9985 | 0.9992 |
| BBMap_fast | 3435.6 | 0.9973 | 0.9989 | 0.9957 |
| BWA_MEM2 | 2142.9 | 0.9718 | 0.9471 | 1.0000 |
| Kraken2Bowtie2 | 142.3 | 0.9986 | 0.9973 | 1.0000 |
| Kraken2HISAT2 | 163.6 | 0.9986 | 0.9973 | 0.9999 |
| Minimap2_illumina | 518.5 | 0.9985 | 0.9970 | 1.0000 |
| Kraken2Minimap2_illumina | 149.1 | 0.9984 | 0.9967 | 1.0000 |
| **Single-end Nanopore** | | | | |
| BioBloom | 346.6 | 0.9755 | 1.0000 | 0.9510 |
| Minimap2_nanopore | 1010.6 | 0.9748 | 0.9996 | 0.9499 |
| KrakenMinimap2_nanopore | 357.3 | 0.9746 | 0.9981 | 0.9510 |
| Kraken2 | 282.7 | 0.9744 | 0.9983 | 0.9504 |

**Table S3 Comparison of the performance of Deconseq and HoCoRT using Bowtie2 in end-to-end mode on single-ended HiSeq and MiSeq reads.** The average runtime (in seconds), accuracy, precision, and sensitivity is shown for each tool. The best (blue) and worst (red) performing tool is indicated for each performance metric.

| Tool and pipeline | Runtime | Accuracy | Precision | Sensitivity |
|---|---|---|---|---|
| **Single-end HiSeq** | | | | |
| Bowtie2_end-to-end | 49.8 | 0.99947 | 0.94934 | 0.99992 |
| Deconseq | 1677.2 | 0.99869 | 0.88432 | 0.99994 |
| **Single-end MiSeq** | | | | |
| Bowtie2_end-to-end | 88.7 | 0.99960 | 0.96194 | 0.99974 |
| Deconseq | 4384.2 | 0.99824 | 0.85028 | 1.00000 |
